# Mapping short-latency cortical responses to electrical stimulation of thalamic motor nuclei by increasing sampling rate - a technical report

**DOI:** 10.1101/788307

**Authors:** Emilia Toth, Ganne Chaitanya, Sandipan Pati

## Abstract

Direct electrical stimulation (DES) of the cortex is a clinically indispensable brain mapping technique that provides reliable information about the distribution of eloquent cortex and its connectivity to the white matter bundles. Here we present a technical report on mapping the short latency cortical responses to stimulation of the ventral anterior nucleus of human thalamus. Reliable downstream responses were noted in the regions connected to the ventral anterior nucleus i.e. superior and inferior frontal gyri, supplementary motor area and limbic substructures (cingulate gyrus and hippocampus).

## Background

Direct electrical stimulation (DES) of the cortex is a clinically indispensable brain mapping technique that provides reliable information about the distribution of eloquent cortex and its connectivity to the white matter bundles(David et al., 2010). Apart from functional mapping, DES is also an efficient way to map neuroanatomical pathways connected to the stimulated node (Duffau, 2015, Lacruz et al., 2007, Martino et al., 2010, Matsumoto et al., 2007). The stimulated node act as an input gate into the large-scale network whose responses are mapped with high temporospatial precision using electrocorticography (ECoG). Evoked neural responses (called as cortico-cortical evoked potentials-CCEP) are measured as the variation in amplitude and first-peak latency (N1) that helps to estimate patient-specific neuroanatomical pathways *in vivo(Duffau, 2015, Keller et al., 2014)*. Evoked cortical responses below 10 milliseconds (ms) reflect mono- or oligosynaptic connectivity while responses longer than 10ms may reflect poly-synaptic connectivity (Keller et al., 2014, Logothetis et al., 2010). Unfortunately, most CCEP studies preclude mapping the early short-latencies (<10ms) as the injected current saturates the amplifier of the clinical EEG acquisition system. Capitalizing on the improved clinical EEG acquisition system (Natus® Quantum®) that allows higher sampling (16 kHz) from 256 channels, we explored mapping short-latency cortical responses (<10ms) to stimulation of ventral anterior nucleus of the thalamus (VA). Based on the known anatomical connectivity of the VA thalamus (often considered a part of the motor thalamus)(Aggleton et al., 1980, McFarland and Haber, 2000, Percheron et al., 1996), we hypothesize that short-latency evoked potentials (SLEP: <10ms) can be mapped in cortices that receive afferents from the VA.

## Methods

The study was performed on a 22-year old right-handed man with drug-resistant focal epilepsy who underwent stereo EEG (SEEG) exploration for localization of seizures. Post-SEEG the seizures were localized to the left orbitofrontal region that was resected, and the patient remained seizure-free (>12 months). One of the depth electrodes sampling the frontal-operculum and insula was progressed medially to sample VA. The study was approved by IRB and a written informed consent was obtained to record field potentials from the thalamus. Stimulation was performed (Nicolet^®^ stimulator) when all antiseizure medications were discontinued to record seizure. Stimulation parameters were: bipolar stimulation (thalamic deepest contact as a cathode and the adjacent contact as anode), 1Hz biphasic pulse wave with pulse width 300 µsec, current 3 mA and train of 40 seconds. The post-implant computed tomography (CT) image was co-registered with the pre-implant structural MRI using Advanced Normalization Tools (Avants et al., 2009). We then combined registration strategies in LeadDBS (Horn and Kuhn, 2015) and iElectrodes (Blenkmann et al., 2017) to map the electrode trajectory and the final thalamic target. The cortical regions implanted were confirmed with AAL2 atlas (Rolls et al., 2015), and the thalamic subnuclei were identified using the mean histological thalamic atlas (Krauth et al., 2010)(Fig. 1K).

**Figure 1:**
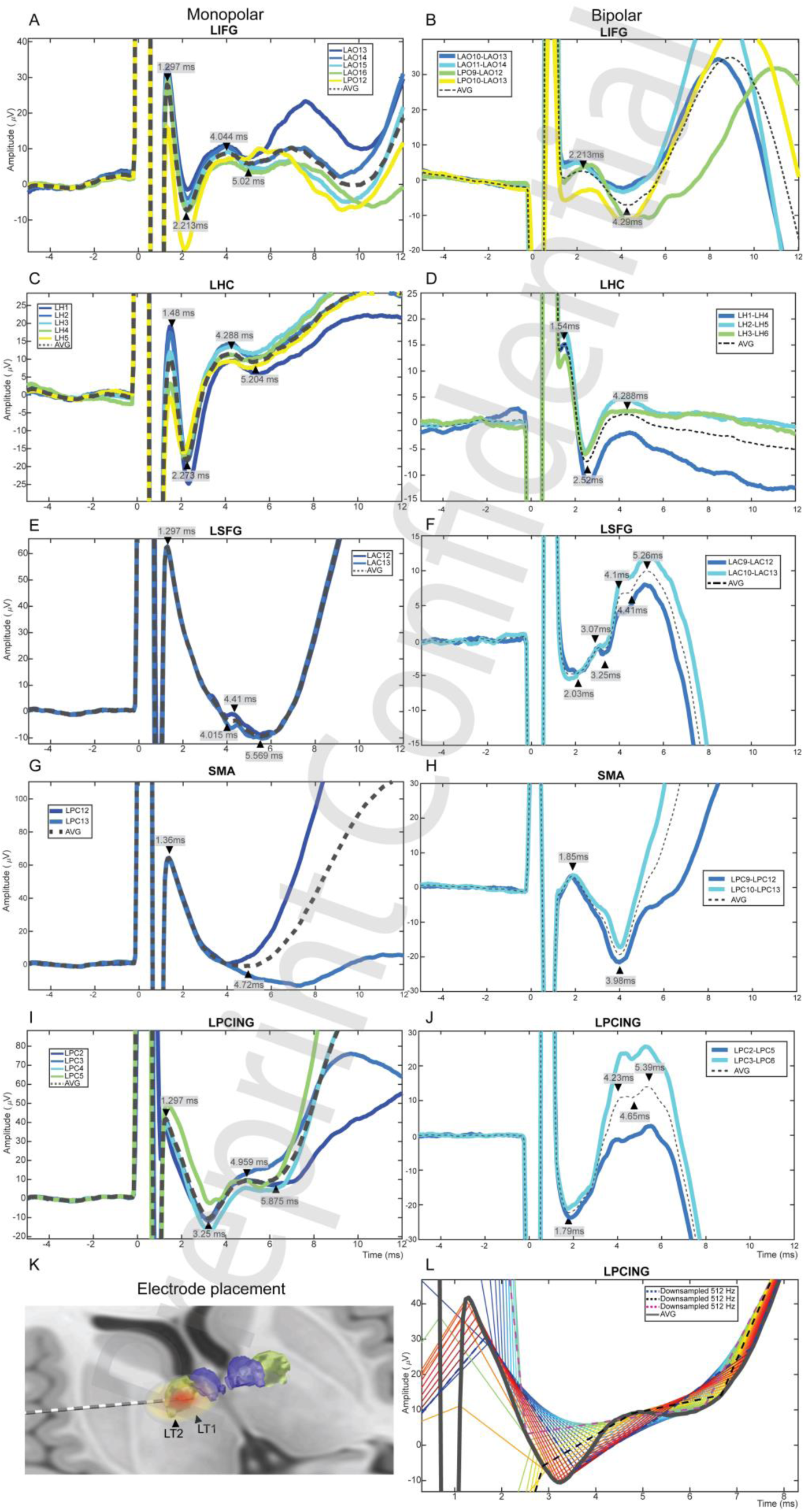
Short latency cortical evoked potentials. Stimulation of the thalamic electrodes elicited early responses that were visible on both monopolar (A, C, E, G, I) and bipolar montages (B, D, F, H, J). With the advantage of a high sampling rate (16KHz), we were able to notice consistent evoked responses at durations varying depending on the cortex sampled. K) Thalamic electrode visualization on patient’s MRI. Thalamic areas VA and ANT signed with green and purple. 3D simulation of the effect and spread of bipolar stimulation of LT1-LT2 channels. L) Different variations (rainbow colored thin lines) of downsampling the average response from Cingulate (magnified from I). All miss the short latency peaks.

## Results

Single pulse electrical stimulation at the left VA thalamus elicited two SLEPs within 10ms that were evident in both unipolar and bipolar montages of SEEG (Fig 1A-J). The SLEPs were localized to left superior and inferior frontal gyrus (LSFG, LIFG), supplementary motor area (LSMA) and limbic substructures (hippocampal gyrus-LHG and posterior cingulate gyrus-LPCING). The earliest response was noted in LIFG and LHG (latencies: 2.21ms and 2.27ms respectively) followed by LSFG and LPCING (latencies: 4.015ms and 3.25ms respectively). Note that the very early peaks of the SLEPs visible on the monopolar (∼ 1.3ms) tended to disappear or shifted (to 1.5-1.8ms) on bipolar montage which implies that the first peak in monopolar is a combination of the fading stimulation artifact and possible cortical response that becomes apparent on the bipolar montage. Sampling rate had a significant effect on detecting these SLEPs. The dotted red and blue lines in Figure 1L demonstrates how down-sampling to 512Hz fails to register the SLEP. In fact, the down-sampled data shows erroneous temporal shift in the detection of the trough.

## Discussion

To date, most CCEP’s were performed with clinical ECoG that were sampled at 500-1000 Hz, and the first 10ms of the evoked response was typically discarded due to uncertainty in distinguishing the neural response from the stimulation-induced artifact (Keller et al., 2014). Here we demonstrate the ability to record SLEP reliably by increasing the sampling frequency with a clinical ECoG acquisition system. Similar SLEPs were localized in the motor cortex with stimulation of the subthalamic nucleus (Miocinovic et al., 2018). Unlike monopolar electrodes that are susceptible to recording of volume conduction from distant sources, bipolar derivation (inter-electrode distance 3.5mm) is likely to represent local sources (Wennberg and Lozano, 2003). Common noise such as the stimulation artifact is likely to have less detrimental effect on the recorded evoked responses. The localization of SLEPs as in Fig 1A-J are consistent with viral tracer studies demonstrating synaptic connectivity of VA with frontal and limbic cortices (Aggleton et al., 1986, Amaral and Cowan, 1980, McFarland and Haber, 2000).

Mapping SLEPs with ECoG is gaining traction, as it identifies variations in neural circuits that can explain inter-individual differences in response to treatment and hence provides the opportunity to individualize therapy (Miocinovic et al., 2018, Walker et al., 2012). For example, SLEPs to pallidal or subthalamic DBS is a candidate biomarker for optimizing DBS target localization and stimulation parameters (Miocinovic et al., 2018). Hence, clinical recording systems need to be innovatively upgraded to effectively map SLEPs reliably in a busy clinical practice without the need for expensive research tools. Analog to Digital Conversion (ADC) forms an essential part of present-day EEG systems that measure continuous analog EEG signal at a discrete-time interval (i.e. sampling at fixed interval) and then digitize the signal. For ADC to accurately reproduce the input complex waveform requires a higher *sampling rate* (measured in hertz) and *resolution* (measured in bits). In Fig1L, we demonstrate the variability in SLEP with down-sampling underscoring the importance of sampling at a higher rate. In summary, we describe the ability to record SLEP from stimulating VA thalamus with a clinical ECoG acquisition system.

